# Genome-wide analysis and evolutionary history of the Necrosis and Ethylene-inducing peptide 1-like protein (NLP) superfamily across the Dothideomycetes class of fungi

**DOI:** 10.1101/2022.02.07.479250

**Authors:** Thaís C. S. Dal’Sasso, Hugo V. S. Rody, Luiz O. Oliveira

## Abstract

Necrosis and Ethylene-inducing peptide 1-like proteins (NLPs) are broadly distributed across bacteria, fungi and oomycetes. Cytotoxic NLPs are usually secreted into the host apoplast where they can induce cell death and trigger plant immune responses in eudicots. To investigate the evolutionary history of the NLPs, we accessed the genomic resources of 79 species from 15 orders of Dothideomycetes. Phylogenetic approaches searched for biased patterns of NLP gene evolution and aimed to provide a phylogenetic framework for the cytotoxic activities of NLPs. Among Dothideomycetes, the NLP superfamily sizes varied, but usually contained from one to six members. Superfamily sizes were higher among pathogenic fungi, with family members that were mostly effector NLPs. Across species, members of the NLP1 family (Type I NLPs) were predominant (84%) over members of the NLP2 family (Type II NLPs). The NLP1 family split into two subfamilies (NLP1.1 and NLP1.2). The NLP1.1 subfamily was broadly distributed across Dothideomycetes. There was strong agreement between the phylogenomics of Dothideomycetes and the phylogenetic tree based on members of the NLP1 subfamilies. To a lesser extent, phylogenomics also agreed with the phylogeny based on members of the NLP2 family. While gene losses seem to have shaped the evolutionary history of NLP2 family, ancient gene duplications followed by descent with modification characterized the NLP1 family. The strongest cytotoxic activities were recorded on NLPs of the NLP1.1 subfamily, suggesting that biased NLP gene retention in this subfamily favored the cytotoxic paralogs.

## INTRODUCTION

Necrosis and Ethylene-inducing peptide 1-like proteins (NLPs) are a superclass of proteins present in bacteria, fungi and oomycetes [1]. In general, NLPs are small proteins of about 24kDa [2] that exhibit cytotoxic activity with cell death-inducing properties and triggers of plant immune response in eudicots, but not in monocots [1–3]. The first purified NLP was the Necrosis- and Ethylene-inducing peptide 1 (Nep1) from *Fusarium oxysporum* f. sp. *erythroxyli*, a protein capable of inducing both necrosis and production of ethylene when applied to *Erythroxylum coca* (coca) [4].

A number of evidence-based studies indicated that NLPs of plant-associated fungi may play a role as virulence factors during the early stages of infection and disease development. The removal of a cytotoxic NLP-encoding gene decreased the virulence of fungal mutants compared to the wild type strain [5]. Conversely, the overexpression of a cytotoxic NLP-encoding gene increased the virulence of the mutant [6]. The role of NLPs in the infection process may not be equally important for all phytopathogens. For example, the removal of NLP-encoding genes did not change the virulence of *Magnaporthe oryzae* [7] and *Botrytis elliptica* [8] mutants. Through functional analyses, some members of the prolific NLP superfamilies of both *Diplodia seriata* [9] and *Neofusicoccum parvum* [10] were shown to exhibit varying levels of cytotoxicity. Contrary to their cytolytic counterpart, the noncytolytic NLPs cannot permeabilize the plant membrane but retain the capability of triggering plant-immune responses; the biological role of the noncytolytic NLPs are yet to be characterized [11–14].

The defining molecular characteristic of an NLP is its NPP1 domain (Pfam PF05630) [15]. This domain contains a highly conserved, seven amino acid long motif (GHRHDWE) that are involved in cation binding [11, 13, 16]. In the N-terminal half of NLPs, there are conserved cysteine residues that are able to form disulfide bridges, which seem to be essential for protein stabilization and necrosis induction [15, 16]. Consistent with their role as proteins of the secretory pathway, the vast majority of NLPs have an N-terminal signal peptide [3, 17].

Based on the number of the conserved cysteines, NLPs have been classified into four main types: a) Type I NLPs have two conserved cysteine residues and are the most widespread type, occurring in fungi, bacteria, and oomycetes [1, 3, 17]; b) A variant called Type Ia also has two cysteines, but differs from Type I by the occurrence of distinct amino acid substitutions; Type Ia is found among oomycetes [17]; c) Type II NLPs have four conserved cysteines, which are responsible for the formation of two disulfide bridges; occurrence of Type II NLPs is scarce, mostly in bacteria and fungi [1]; and d) Type III NLPs exhibit the least conserved amino acid sequence, most of them carry six cysteine residues that are able to form three disulfide bridges; they occur in ascomycetes and some bacteria [3]. The occurrence of Types I, II, and III NLPs in Ascomycota suggests that the NLPs originated within this phylum; horizontal gene transfer (HGT) is a plausible mechanism that may have allowed NLPs to reach distantly-related taxa [17].

A recent study surveyed the taxonomic distribution of NLPs in reference proteomes of about 10 thousand species of bacteria, fungi, and oomycetes; about 500 species contained at least one NLP [3]. The most striking features that bring together most of the species that do contain NLPs are: their taxonomic placement within the phylum Ascomycota (60%, 211 of 360 proteomes), their plant pathogenicity (some of the most obnoxious phytopathogens do contain NLPs), and their trophic modes (especially pathogenic and saprophytic microbes). In Dothideomycetes, the overall number of NLPs copies is much smaller than the extremely large copy size number found in Oomycetes, in which up to 100 copies have been documented in some genomes [3]. Undoubtedly, the diversification of the NLPs across the Dothideomycetes is a complex process, and far from being understood.

The Dothideomycetes comprises the largest and phylogenetically most-diverse class within the phylum Ascomycota, with an estimated number of members that may reach up to 19 thousand species [18, 19]. They occur across diverse habitats, including extreme environmental conditions, and their lifestyles are very diverse [18, 20, 21]. Dothideomycetes diverged from other sister Ascomycota classes around 366 million years ago (mya) [22]. Currently, the class comprises 23 orders [20] that are believed to have evolved in the range between 100 and 220 mya [22]. The Pleosporales, the Capnodiales, and the Botryosphaeriales are three of the largest orders within the Dothideomycetes; each of those orders contains a large number of families, which in turn hold some of the most destructive genera of plant pathogens to cereal crops, trees, and dicots [20, 21, 23].

With incredible ecological and morphological diversities and economical importance as plant pathogens, the Dothideomycetes have raised the interest of genomic research in the recent decade. Currently, genomes of about 90 genera of Dothideomycetes are available in public repositories, most of which are from plant pathogens and plant-associated species [20, 23]. Those genomic resources allow for the investigation of the evolutionary history of NLPs across that class.

We began our study by building up a robust molecular phylogenetic framework based on a set of 1,851 single copy ortholog (SCO) proteins from 79 species of Dothideomycetes. To account for ecological diversity within the class, we included species of three trophic modes (pathotrophs, saprotrophs, and symbitrophs) sorted out among plant pathogens, plant-associated species, and non-plant associated species. Built independently from the NLP evolutionary history, our framework revealed the phylogenomic relationships among the studied species, and should shed light into the evolutionary history of NLPs across Dothideomycetes.

Next, we reconstructed the phylogenetic relationships among NLPs (Types I, II, or III) based on the sequences we had retrieved from those genomic resources and questioned how ubiquitous and diverse the NLP superfamily became across the Dothideomycetes. If NLP gene evolution took place in an NLP type-independent manner, the three types of NLPs should show similar distribution across the phylogenies. Otherwise, if biased patterns drove NLP gene evolution, the NLP-based phylogenies will be imbalanced owing to unequal gene losses, gene duplications, and gene retention that took place over time. As a consequence, NLP types that experienced adverse selection will be rare; likely vanishing from certain subclades.

Subsequently, we explored how the processes that drove NLP gene evolution took place at the lower ranks of the NLP phylogenetic hierarchy. To answer our questions, we build independent phylogenies, one for each of the major NLP subclades we had recovered previously. Then, we contrasted the topology of those trees with the phylogenetic framework of Dothideomycetes.

Finally, we provide a context to explore whether the cytotoxic activities reported previously through functional analyses are associated with NLP gene evolution and phylogenetic relationships among species. If preferential gene retention patterns maintained functional paralogs over time, cytotoxic activities will tend to exhibit some phylogenetic signals. This investigation shed light on the likely mechanisms that contributed to the evolution of NLPs in Dothideomycetes.

## MATERIAL AND METHODS

### Data assembly and trophic mode prediction

From MycoCosm [24], we retrieved protein sequences of 79 species (representing 15 orders) of Dothideomycetes. There was one species per genus, and two additional species of Eurotiomycetes (*Aspergillus fumigatus* and *Aspergillus nidulans*). The 81 species (79 Dothideomycetes and two *Aspergillus* spp.) used in this study are listed (Additional File 1)

To infer the trophic modes, we run the FUNGuild annotation tool locally [25] using the database for fungi and a list of species (Additional File 1) as input.

### Orthogroups and phylogenomic analysis

OrthoFinder v2.3.3 [26] established orthologous relationships among members of the Dothideomycetes (Additional File 1). OrthoFinder calculated length and phylogenetic distance-normalized scores from an all-versus-all alignment using DIAMOND v0.9.24 [27] and identified reciprocal best normalized hits. Normalized scores above default thresholds were assigned to the orthogroup graph and subjected to the Markov clustering analysis in order to assume ortholog groups (orthogroups).

We used a Maximum-likelihood phylogeny approach to uncover the phylogenetic relationships among species of the Dothideomycetes. We firstly prepared a dataset with the SCO proteins as identified by OrthoFinder. Proteins were aligned using the L-INS-I method as implemented in MAFFT v7.453 [28]. TrimAL v1.4.rev22 [29] trimmed the alignments, with the parameters “-gt 0.95”, and “-cons 60”. After trimming and concatenation, we obtained dataset 1.

We subjected dataset 1 as input to IQ-TREE v1.6.11 [30] to estimate both the best evolutionary model and the phylogenomic tree. According to the Bayesian information criteria (BIC), IQ-TREE standard model selection indicated LG+F+I+G4 as the best fit model. Subsequently, we carried out a Maximum-likelihood phylogenomic tree using 1,000 ultrafast bootstrap replicates and ten independent runs. *Aspergillus fumigatus and Aspergillus nidulans* were set as outgroups. The tree was visualized in FigTree v1.4.4 (http://tree.bio.ed.ac.uk/software/figtree/).

### Protein annotation and assemble of NLP homologues

The obtained proteomes of the set of 81 species were annotated using PfamScan with Pfam v32.0 [31] and InterProScan v5.30.69 [32] with the following eight parameters: SMART-7.1, SUPERFAMILY-1.75, ProDom-2006.1, CDD-3.16, TIGRFAM-15.0, Pfam v31.0, Coils-2.2.1, and Gene3D-4.2.0.

The presence of a signal peptide (according to SignalP v4.1; [33]) and the absence of transmembrane domains (according to TMHMM v2.0; [34]) defined the proteins that we predicted to be part of the secretome. Subsequently, TargetP [35] predicted the subcellular localization of the predicted secretome.

Finally, HMMER v3.2.1 (http://hmmer.org) predicted the NLP homologues (E-value < 0.001) using profile hidden Markov models (HMMs) for NPP1 domain (PF05630) from the proteomes. A protein was declared to be an NLP when the NPP1 domain was annotated by at least two out three softwares: PfamScan, InterproScan, and HMMER. Within the pool of NLPs, a given protein was considered to be an ‘effector NLP’ when it passed the following three tests: (a) SignalP predicted it harbors a signal peptide, (b) TMHMM predicted it has no transmembrane domain, and (c) TargetP predicted it to be part of a secretory pathway. When a given NLP failed any of these three tests, it was regarded as a ‘non-effector NLP’. Finally, we assembled dataset 2, which contained the predicted NLPs (protein data) that were present in the set of 81 species.

### NLP Maximum-likelihood phylogenies

Maximum-likelihood phylogenetic analysis based on dataset 2 allowed us to reconstruct the phylogenetic relationships of NLPs among the members of the Dothideomycetes.

Alignments were obtained using the L-INS-I method, as implemented in MAFFT v7.453. MAFFT L-INS-I allows for aligning a set of sequences containing sequences flanking around one alignable domain with high accuracy [28]. To find the best fit model, we used dataset 2 and its partitions as input to IQ-TREE. According to BIC, IQ-TREE selected WAG+F+I+G4 for dataset 2; for its partitions, the best evolutionary models were WAG+F+I+G4, WAG+I+G4, or VT+G4. Maximum-likelihood phylogenetic trees for NLPs were also performed in IQ-TREE as described previously. Phylogenetic analysis of the full dataset 2 was performed without outgroup. However, phylogenetic analyses on partitions of dataset 2 (see results section) were carried out using *A. fumigatus* as outgroup.

### Survey on functional activities of NLPs

We surveyed the literature for functional analyses of NLPs across Ascomycota. We recorded the taxonomic placement of the study species, the NLP names attributed to the proteins, and the original outcome of the functional analyses (usually each study ranked the activity as either strong, weak, or absent). We included in our phylogenetic approach the Dothideomycetes species for which a functional analysis had been carried out. Species of other classes of fungi were not included in our phylogenies, but we carried out protein sequence alignments individually in order to assign the placement of each of their NLPs into the phylogenetic treatments we developed herein.

## RESULTS

### Genome-wide phylogeny of Dothideomycetes

The set of 79 species (79 genera, 15 orders) of Dothideomycetes allowed OrthoFinder to uncover 1,851 SCO proteins (a concatenated set of 910,900 amino acids). The whole-genome data provided support for the phylogenetic reconstruction within Dothideomycetes (Fig. 1). The phylogenomic tree showed well-supported nodes (bootstrap values = 100, for all nodes). Overall, there were two major clades. The first major clade held 17 species of four orders (Capnodiales, Myriangiales, Dothideales, and Trypetheliales). The second major clade held 62 species within 11 orders of Dothideomycetes; it was split further into two sister sub-clades. The first of these sub-clades encompassed seven species of four orders: Aulographales (two species), Venturiales (three), Mycrothyriales (one), and Eremomycetales (one). The second sub-clade comprised 55 species of seven orders: Pleosporales (38 species), Hysteriales (two), Mytilinidiales (five), Acrospermales (one), Botryosphaeriales (seven), Patellariales (one), Lineolatales (one).

**Fig. 1.**
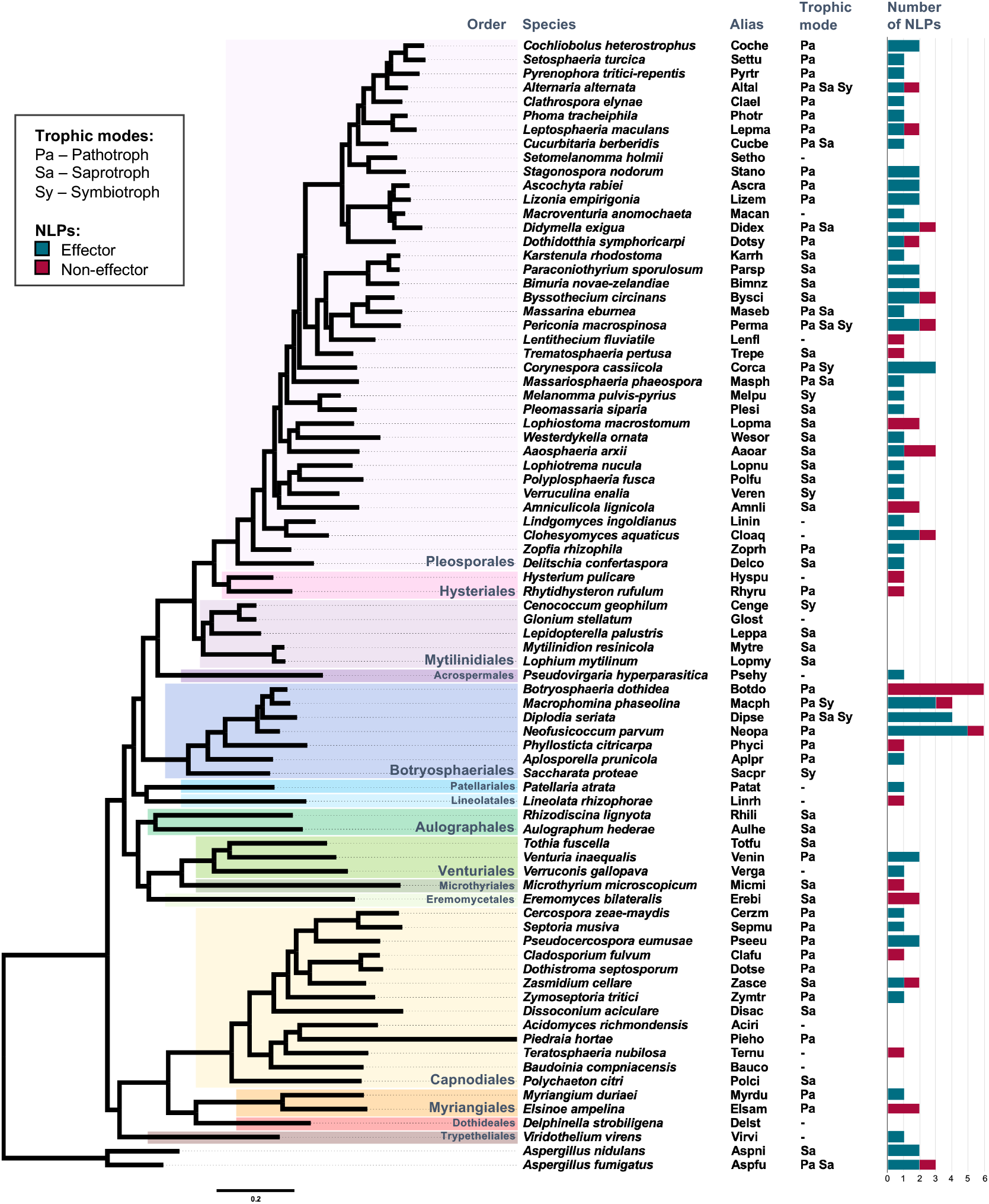
Phylogenomic relationships among 79 species of Dothideomycetes. The Maximum-likelihood consensus tree is based on a dataset of 1,851 single copy ortholog proteins, with *Aspergillus* spp. (Eurotiomycetes) as outgroups. Clades are color-coded according to the orders of Dothideomycetes, as indicated. Five-letter codes indicate alias of each species name. Trophic modes (according to FUNGuild classification) are coded as indicated. Bars indicate the number of NLPs (effectors and non-effectors) per species. Branch lengths are drawn to scale; nodal support values are equal to 100 local bootstraps for all nodes. Scale bar corresponds to the expected number of substitutions per site.

### Genome-wide identification of homologues of NLPs in Dothideomycetes

Apart from the NPP1 domain (PF05630), our pipeline predicted no other domain as part of the NLPs. Each NLP had a single copy of the NPP1 domain. Among the NLPs, our pipeline distinguished between two sets of proteins: (a) effector NLPs and (b) non-effector NLPs. In the first set, the protein harbored a signal peptide, had no transmembrane domain, and was part of the secretory pathway. In the second set, despite the presence of the NPP1 domain, the protein failed to comply with any of the previous three requirements.

In general, NLP superfamily sizes (effector NLPs + non-effector NLPs) among species of Dothideomycetes varied (Fig. 1, Additional File 1). There were 17 species without NLPs: six species of Capnodiales (*Acidomyces richmondensis, Baudoinia compniacensis, Dissoconium aciculare, Dothistroma septosporum, Piedraia hortae*, and *Polychaeton citri*), all five species of Mytilinidiales (*Cenococcum geophilum, Glonium stellatum, Lepidopterella palustris, Lophium mytilinum*, and *Mytilinidion resinicola*), the two species of Aulographales (*Aulographum hederae* and *Rhizodiscina lignyota*), one species of Pleosporales (*Setomelanomma holmii*), one of Botryosphaeriales (*Saccharata proteae*), one of Dothideales (*Delphinella strobiligena*), and one of Venturiales (*Tothia fuscella*).

Among the species that do harbor NLPs, the copy number varied from one (62 species) to six (*Botryosphaeria dothidea* and *Neofusicoccum parvum*; Botryosphaeriales) (Additional File 1). In a given species, the general trend was that the number of effector NLPs was larger than the number of non-effector NLPs. The copy number of effector NLPs ranged from one (32 species) to five (*Neofusicoccum parvum*). In 14 species, however, the NLP family contained exclusively non-effector NLPs (from one to six members).

### Broad sense phylogeny of NLPs

After carrying out a phylogenomic reconstruction within the set of 79 species of Dothideomycetes (Fig. 1), we reexamined the phylogenetic relationships among species with protein data from the NLPs as the only source of information. Therefore, 17 species that lacked NLPs did not take part in this new analysis. The Maximum-likelihood phylogenetic tree contained a total of 111 NLPs, 106 of them came from a total of 62 species of Dothideomycetes; five NLPs came from *Aspergillus* spp. (Fig. 2, Additional File 2). The phylogenetic tree showed well-supported nodes (bootstrap values > 95, for the main nodes). There were three major clades. A shared trait of all NLPs within a given major clade was the number of conserved cysteine residues in the protein (which separates NLPs into Types: I, II, or III). Notably, the placement of non-effector NLPs was spread across the phylogeny.

**Fig. 2.**
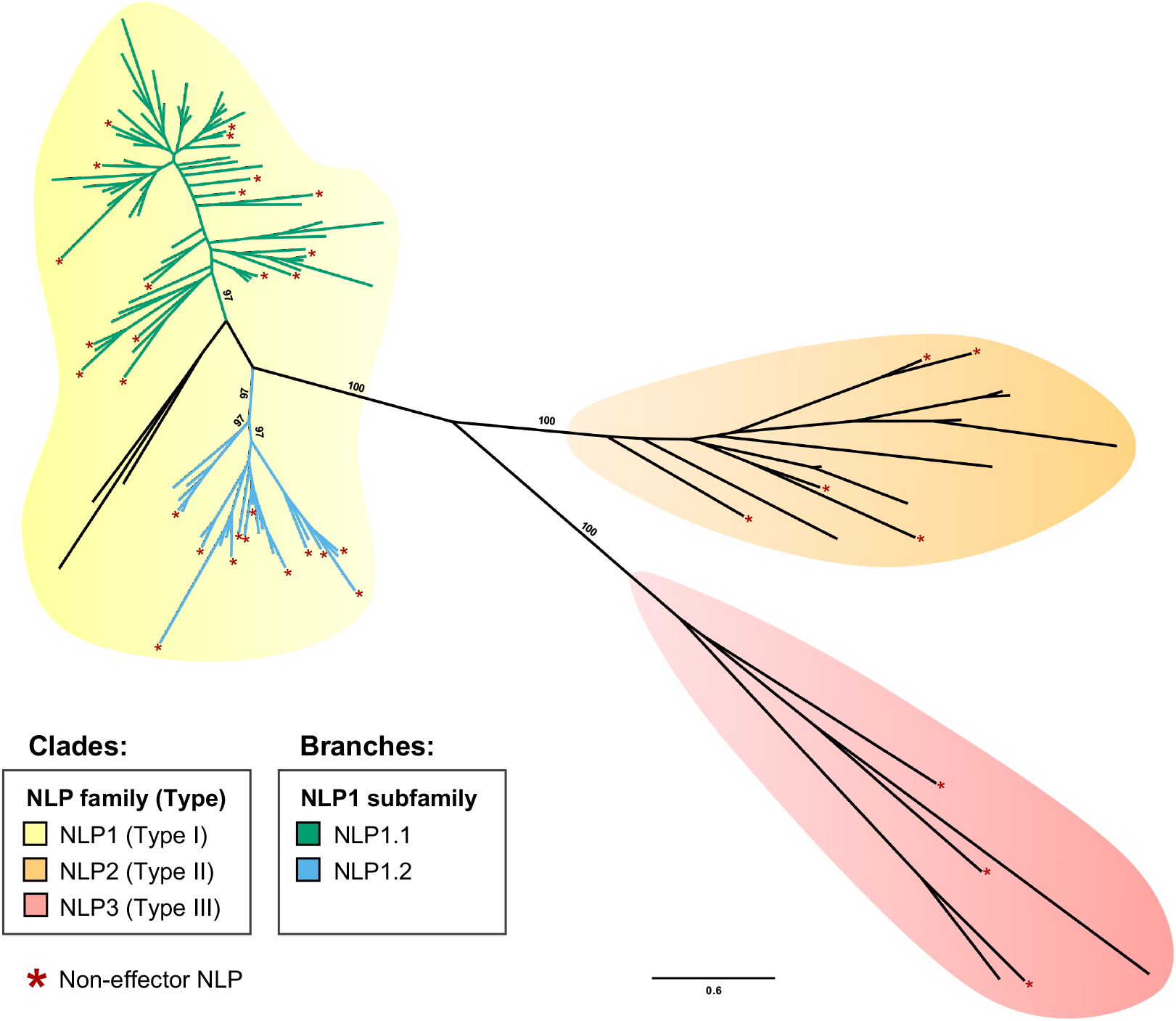
Phylogenetic relationships among 111 predicted Necrosis- and Ethylene-inducing like proteins (NLPs). The dataset for the unrooted Maximum-likelihood phylogeny (consensus tree) comprised proteins that were 572 amino acids long. A total of 106 out of 111 NLPs were from the Dothideomycetes. Clades are color-coded according to the number of conserved cysteine residues in the protein (Type I, two residues; Type II, four residues; Type III, variable number of residues), as indicated. Branches are color-coded according to the NLP1 subfamily, as indicated. Along the phylogeny, asterisks indicate the phylogenetic placements of non-effector NLPs. Branch lengths are drawn to scale; nodal support values are given as local bootstraps (when ≥ 95, for the main nodes) above the branches. Scale bar corresponds to the expected number of substitutions per site.

The first major clade harbored 93 NLPs, all of which were considered as Type I NLPs (with two cysteine residues; see [3]). Among the 93 NLPs, 75 (81%) were predicted to be effector NLPs. The first major clade was split further into three sub-clades. One of the three sub-clades contained three highly differentiated effector NLPs: one from *Aaosphaeria arxii* (Aaoar_355692), one from *Paraconiothyrium sporulosum* (Parsp_876375), and one from *Periconia macrospinosa* (Perma_691703). Species composition among the remaining two sub-clades was large and very diverse. The second major clade brought together 13 NLPs (Type II NLPs; with four cysteine residues). Among the 13 NLPs, eight (62%) were predicted to be effector NLPs. Meanwhile, the third major clade encompassed only five NLPs (Type III NLPs; with variable number of cysteine residues). Among them, only two (40%) were predicted to be effector NLPs.

### Narrow sense phylogeny of NLPs

We carried out four subsequent studies based on the phylogenetic relationships we recovered previously (Fig. 2, Additional File 2). The data from the first major clade gave rise to two independent phylogenies (one for each of the two best-resolved sub-clades (Fig. 3, Additional File 2). Hereafter, these two narrow sense phylogenies are referred to as ‘NLP1.1’ and ‘NLP1.2’ phylogenies, respectively. We also built independent phylogenies for the remaining two major clades (Fig. 4, Additional File 2). Hereafter, these two narrow sense phylogenies are referred to as ‘NLP2’ and ‘NLP3’ phylogenies, respectively.

**Fig. 3.**
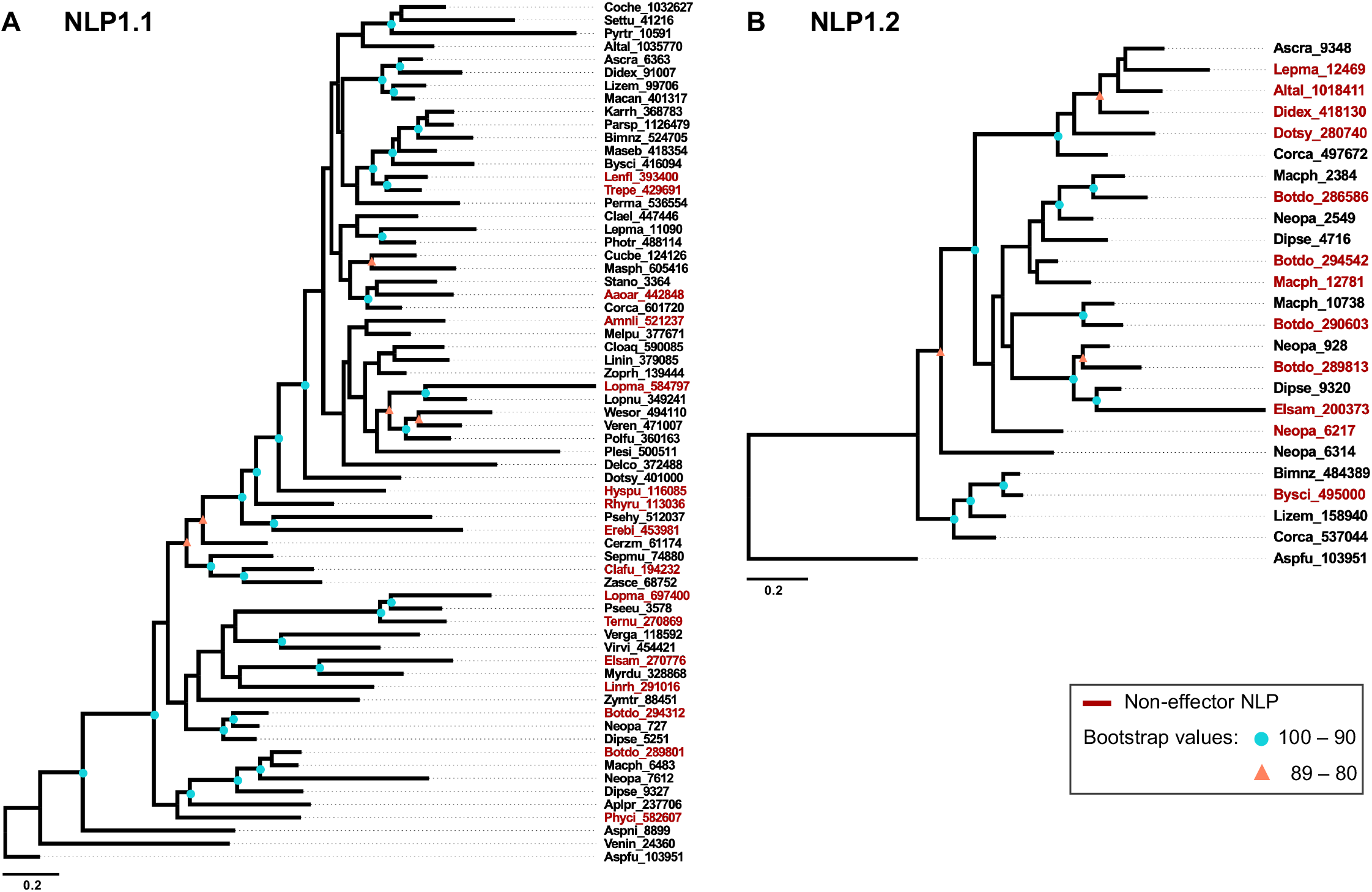
Phylogenetic relationships among members in two subfamilies of the Necrosis- and Ethylene-inducing like protein family 1 (NLP1) in Dothideomycetes. Each phylogeny represents a subclade of NLP1 family depicted in Fig. 2: **A.** Members of NLP1.1 (66 proteins that were 307 amino acids long); **B.** Members of NLP1.2 (24 proteins that were 270 amino acids long). Protein IDs in red indicate non-effector NLPs. For each phylogeny, Aspfu_103951 (NLP1.1) from *Aspergillus fumigatus* (Eurotiomycetes) was used as outgroup. Branch lengths are drawn to scale; nodal support values are given as local bootstraps, as indicated. Scale bar corresponds to the expected number of substitutions per site.

**Fig. 4.**
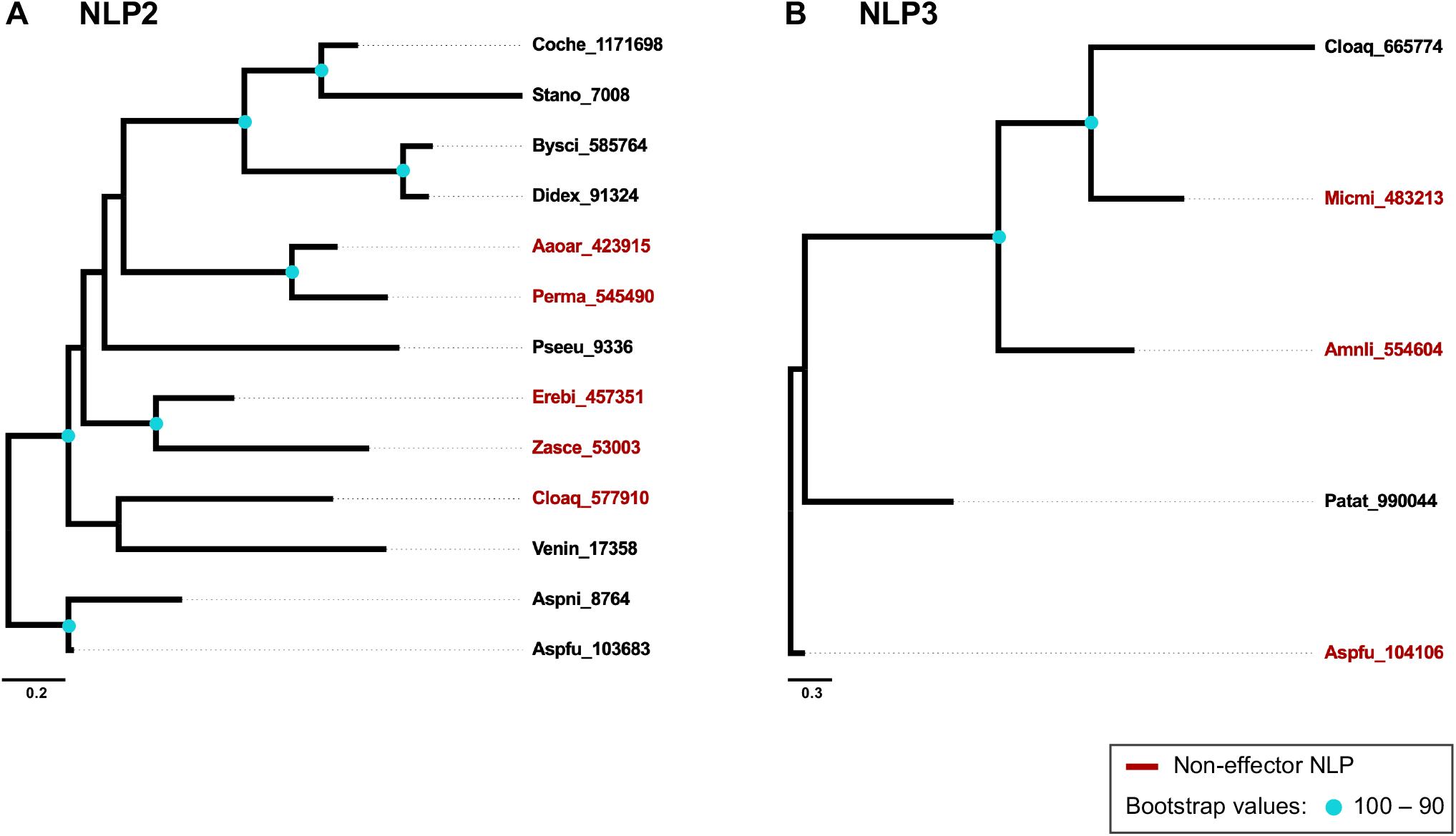
Phylogenetic relationships among members of two Necrosis- and Ethylene-inducing like protein families – NLP2 and NLP3 – in Dothideomycetes. Each phylogeny represents a subclade depicted in Fig. 2: **A.** Members of NLP2 (13 proteins that were 491 amino acids long); **B.** Members of NLP3 (five proteins that were 390 amino acids long). Protein IDs in red indicate non-effector NLPs. For each phylogeny, an NLP from *Aspergillus fumigatus* (Eurotiomycetes) was used as outgroup. Branch lengths are drawn to scale; nodal support values are given as local bootstraps, as indicated. Scale bar corresponds to the expected number of substitutions per site.

Unequivocally, NLP1.1 was the most member-rich phylogeny; it contained 66 Type I NLPs (50 effectors, 76%) from 60 species of Dothideomycetes and *Aspergillus* spp. (Fig. 3A). Amongst the 37 species that contained a single NLP, 34 (92%) of them were present in NLP1.1. Among the 64 NLP1.1 proteins from Dothideomycetes, there were a total of 48 effector NLPs. The richness towards effector NLPs was higher among Pleosporales, in which 32 out of 38 NLPs were predicted as effector proteins.

The NLP1.1 phylogeny split further into two major clades (Fig. 3a). The major clade 1 contains six NLPs from Botryosphaeriales (Botdo_289801, Macph_6483, Neopa_7612, Dipse_9327, Aplpr_237706, Phyci_582607). The major clade 2 was member-richer and split further into two sister sub-clades; the smallest of which contained 12 NLPs from seven orders (Botryosphaeriales, Pleosporales, Lineolatales, Venturiales, Capnodiales, Myriangiales, and Trypetheliales). Finally, the large subclade 2 contained NLPs from Capnodiales, Eremomycetales, Acrospermales, Hysteriales, and a large second-order sub-clade of NLPs from Pleosporales. Overall, NLP1.1 paralleled quite well the molecular phylogenetic framework of Dothideomycetes, especially across Pleosporales. Botryosphaeriales retained a major gene duplication of NLP1.1, with sets of corresponding paralogs in distinct genera being retained over time. This major gene duplication is absent from either Capnodiales or Pleosporales.

The remaining 24 Type I NLPs from fourteen species of Dothideomycetes were analyzed in the NLP1.2 phylogeny; half of the members (12 out 24) were non-effector NLPs (Fig. 3b). This phylogeny revealed two additional duplication events; one of them was shared among Botryosphaeriales only. The second duplication gave rise to a sub-clade that has a sister relationship to Pleosporales. Although not as member-rich as NLP1.1, NLP1.2 also resembled well the molecular phylogenetic framework of Dothideomycetes.

The Type II NLPs clustered together in the NLP2 phylogeny; they were of rare occurrence (nine out of 79 species) amongst the Dothideomycetes and *Aspergillus* spp.; five species were Pleosporales (Fig. 4a). Finally, there was the NLP3 phylogeny, which consisted of only five Type III NLPs, from four species amongst the Dothideomycetes and *Aspergillus fumigatus* (Fig. 4b).

### Cytotoxic NLPs within a phylogenetic context

Our survey recovered 27 functional analyses of NLPs (Table 1). There were 11 instances in which the authors attributed to the NLP under investigation a strong activity, all of the proteins were Type I NLPs. Among those 11 proteins with strong cytotoxic activity, there were seven NLP1.1 and four NLP1.2. There were nine instances in which a Type 1 NLP showed weak/absent activity; those instances took place in species with multiple copies of the Type I NLPs and a strong activity already had been attributed to one of the NLP1 paralogs. Finally, all six functional analyses of Type II NLPs (NLP2) and a single analysis of Type III NLP (NLP3) pointed to weak/absent activity. In *Neofusicoccum parvum* (Botryosphaeriales), the NLP1.2 paralogs Neopa_928 and Neopa_6217 had been both removed from functional analysis experiment [10]. While the overexpression of Neopa_928 as a recombinant protein was not possible; Neopa_6217 coded for a truncated protein that lacked signal peptide. Meanwhile, our analyses predicted that Neopa_928 was an effector NLP and Neopa_6217 was a non-effector NLP.

**Table 1.**
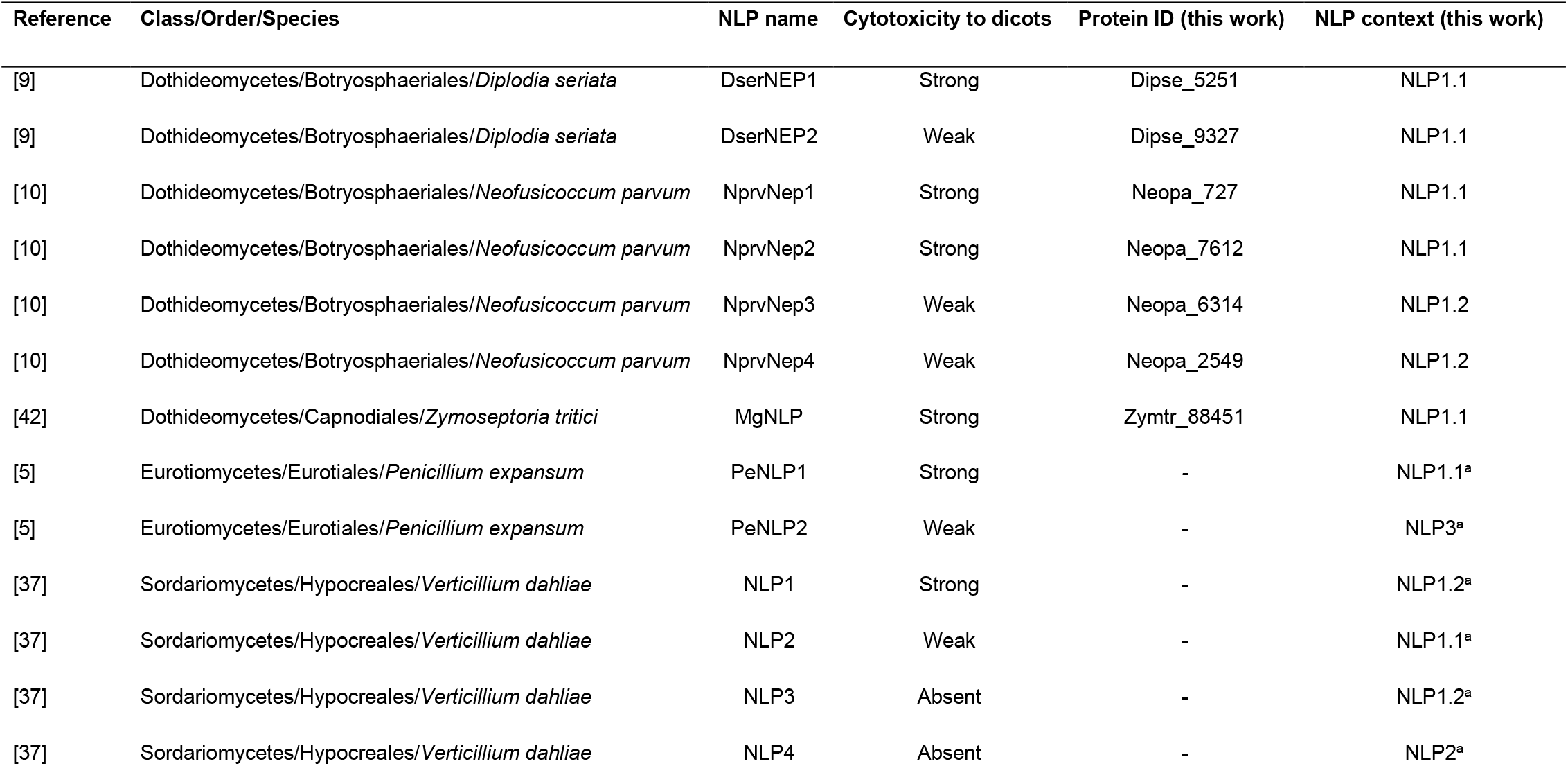
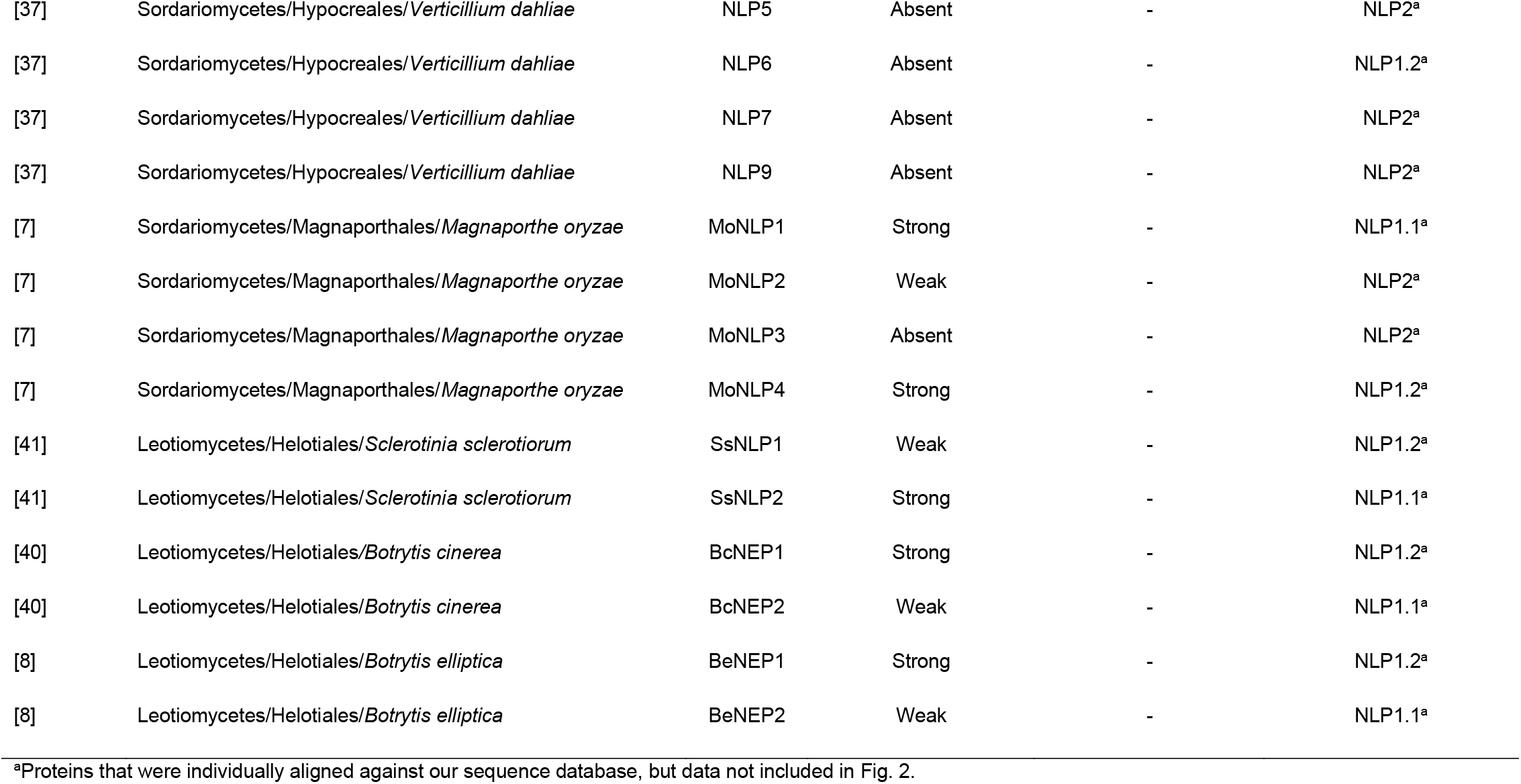
The cytotoxicity of NLPs found in species of the phylum Ascomycota.

### NLPs within a trophic mode context

Using the FUNGuild annotation tool, we inferred the trophic mode for each of the Dothideomycetes species into three groups: pathotrophs, saprotrophs, and symbiotrophs (Fig. 5, Additional File 1). Sixty-three (80%) out of 79 species of Dothideomycetes were classified into at least one of the three groups. There were 26 pathotrophs, 25 saprotrophs, and four symbiotrophs. Eight species were classified as part of more than one group: pathotrophs-symbiotrophs (two species), pathotrophs-saprotrophs-symbiotrophs (three), and pathotrophs-saprotrophs (three). In general, the average number of NLPs and effector NLPs per species was higher amongst the species classified as pathotrophs (exclusively or not).

**Fig. 5.**
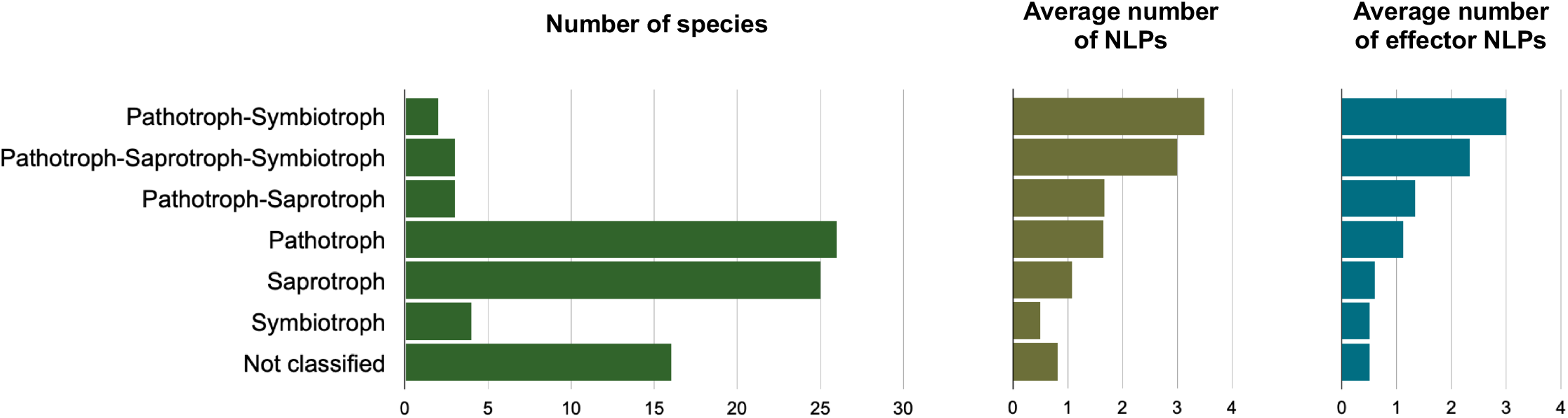
Trophic modes predicted among 79 species of Dothideomycetes and its relationship with NLP copy number. The total number of species per trophic mode is shown on the left chart. The average numbers of NLPs and effector NLPs per trophic mode per species are shown on the central and right charts, respectively. Classification of Dothideomycetes into trophic modes was performed using the FUNGuild annotation tool.

## DISCUSSION

### The NLP superfamily

The current type-based classification of NLPs (Types I, II, and III) relies exclusively on the number of conserved cysteine residues that form disulfide bridges along the N-terminal half of the proteins [1, 17]. Previously, a broad phylogenetic analysis of NLPs from bacteria, fungi and oomycetes [3] provided hints that suggested NLPs of a given type might share a common ancestor. Herein, we used phylogenomics and NLP phylogenetic data to explore the evolutionary history of NLPs in the Dothideomycetes class of fungi.

Our studies support a phylogeny-based classification of NLPs into a superfamily. Accordingly, our phylogenetic framework uncovered at least two subordinate families that corresponded to existing NLP types. Hereafter, we will refer to Type I NLPs collectively as the ‘NLP1 family’, and Type II NLPs as the ‘NLP2 family’. Furthermore, the phylogenetic relationships within the member-rich NLP1 family of Dothideomycetes supported two further natural subdivisions: hereafter referred to as NLP1.1 and NLP1.2 subfamilies.

Likely, Type III NLPs may also merit their own phylogenetic-based category into a NLP3 family; however, several aspects of their evolutionary history are still unclear, such as their low sequence conservation and likely association with horizontal gene transfer [17]. Moreover, the rare occurrence of Type III NLPs in Dothideomycetes may bias our studies. Thus, we refrained from making further attempts to explore the phylogenetics of NLP3 family with the current dataset available to this investigation.

The NLP superfamily exhibited a small size across Dothideomycetes, with one (62 species) to six members (two species in Botryosphaeriales). Among Dothideomycetes, NLP family size is likely associated with trophic mode. Fungi inferred to be pathotrophs detained the highest number of NLP copies. Cell death-inducing proteins, such as NLPs, which act at the plant apoplast level, are essential for host colonization [9, 10, 36]. Moreover, the role of the larger NLP family in necrotrophic plant pathogens, such as that of the Botryosphaeriales, can go beyond differential cytotoxicity and detain levels of functional diversification at different life stages [37, 38] or, to some extent, contribute to infection of wider host ranges by those pathogens [37, 39].

### Biased patterns of gene evolution drove NLP diversification

The repositories of NLPs are the product of dynamic events that took place during the evolutionary history of extant Dothideomycetes. The NLP1 family and the NLP2 family clearly underwent distinct pathways during their evolution. Notably, gene retention and gene loss over time are major drivers that distinguish the NLP1 family apart from NLP2 family.

The NLP1 family likely underwent rounds of gene duplication followed by descent with modification, which allowed for its diversification into at least two subfamilies (NLP1.1 and NLP1.2). However, these subfamilies experienced distinct evolutionary processes over time. Preferential gene retention contributed to preserve the NLP1.1 paralogs that emerged after gene duplication and explains the ubiquitous presence of the NLP1.1 subfamily members across most of the extant Dothideomycetes (Fig. 3a). On the contrary, gene loss is a realistic scenario that explains the evolutionary history of the NLP1.2 subfamily. Over time, preferential gene loss eliminated NLP1.2 members from the genomes of most extant Dothideomycetes (Fig. 3b). The fact that the phylogenies of each subfamily resembled the phylogenomics of their species supports the scenario of early gene duplications and subsequent biased gene losses in some lineages as a likely explanation to account for the extant pattern of distribution of the NLP1 family across Dothideomycetes.

The signatures of past lineage-specific gene duplication and gene retention are visible in the phylogenies of subfamilies NLP1.1 and NLP1.2, especially in the order Botryosphaeriales, which showed above-average number of the NLP1 family (Fig. 3). We speculate that the otherwise redundancy of extant copies in Botryosphaeriales may contribute to shape a strategy of functional diversification to overcome host-related defense mechanisms, with distinct NLPs playing a functional role in a time-dependent manner to allow successful colonization of plant tissues.

In sharp contrast to the ubiquitous presence of NLP1 family members, the occurrence of the NLP2 family members among Dothideomycetes was rare. Similarly, to the NLP1.2 subfamily, biased gene losses also seem to have played a major role in driving the trajectory of the NLP2 family. Yet, few members of the NLP2 family survived and are distributed across phylogeny of the extant Dothideomycetes (Fig. 4a).

Non-effector NLPs were those NLPs that do harbor NPP1 domain (PF05630) but either do not detain a signal peptide, may contain transmembrane domains or are not part of the secretory pathway. Apparently, a non-effector NLP lacks some of the key traits that define a protein as a functional NLP. Their presence across the phylogeny implies that non-effector NLPs originated multiple times over time. Along their evolution, likely, the proteins we predicted to be non-effector NLPs underwent independent rounds of sequence rearrangements, with losses and gain of traits, including their secretion signals. High frequency of non-effectors among NLP1.2 paralogs also suggests that ongoing pseudogenization may be helping to avoid functional redundancy of NLPs in Dothideomycetes. The retention of those genes over time is puzzling; whether this is a strategy to escape from host recognition remains to be demonstrated experimentally.

### Phylogenetic framework of NLP cytotoxicity

Through functional analyses, a number of NLPs have been demonstrated to exhibit cytotoxic activity in eudicot plants [5, 7–10, 37, 40–42]. These analyses were carried out at the species level; the authors originally summarized the final outcomes of the cytotoxicity levels of a given NLP as either ‘strong’, ‘weak’, or ‘absent’. In the present investigation, we demonstrated that the cytotoxicity of NLPs detains a strong phylogenetic signal (Table 1).

We applied our phylogenetic framework to those NLPs that had undergone functional analyses in previous studies. The vast majority of the NLPs that had exhibited ‘strong’ activity were phylogenetically related; they belong to the member-rich family NLP1 and grouped mostly within subfamily NLP1.1. Our findings suggested that members of the NLP1 family may have been favored through biased gene retention over time. Likely, the NLP1 paralogs that encode for proteins with ‘strong’ cytotoxicity are actually encoding effector NLPs with a functional role in promoting virulence. Therefore, the maintenance of these effector-encoding genes in the genome would be advantageous at evolutionary time scale and could explain their maintenance across extant Dothideomycetes.

Experiment-based evidence supports NLP1 members as genes that encode for essential virulence factors in plant pathogenic fungi. In the postharvest pathogen *Penicillium expansum* (Eurotiomycetes), the deletion *in vitro* of a member of the NLP1 family led to the reduction of virulence; meanwhile the virulence remained unchanged when a Type III NLP was the deleted copy [5]. In *Neofusicoccum parvum* (Dothideomycetes), NLP1.1 paralogs encoded proteins that were strongly cytotoxic to both tomato plants and mammalian cells, while NLP1.2 paralogs encoded for proteins that exhibited weak cytotoxicity [10]. The only NLP from *Zymoseptoria tritici* (Dothideomycetes), which induced defense responses and cell death in dicots but not in monocots [42], shared homology to members of the NLP1.1 subfamily. Paralogs of the NLP1.1 subfamily of *Diplodia seriata* (Dothideomycetes) encoded proteins that showed distinct levels of cytotoxicity to grapevine leaves [9].

Our phylogenetic framework grouped most of the paralogs that encode for NLP proteins with ‘weak’ cytotoxicity — as well as proteins with activity classified as ‘absent’ — together with members of either the NLP2 or NLP3 families. The scarcity of either NLP2 or NLP3 members in the genome of most species of Dothideomycetes suggests that the proteins these genes encode for likely play a secondary role in virulence. In some instances, NLP cytotoxicity and immunity induction are uncoupled, and non-cytotoxic NLPs may retain the ability to trigger host immune responses [11, 13]. An additional evidence for a secondary role of NLP2 and NLP3 family members in virulence comes from the fact that fungal genomes that do possess NLP2 or NLP3 genes frequently harbored at least one NLP1 gene that encoded for a protein with ‘strong’ activity in the functional analyses. Therefore, these functional studies support an evolutionary scenario in which members of the NLP1 family likely play a critical role as effector genes; therefore, contributing actively as a tool to overcome host defense systems and facilitate the infection process.

### Conclusion

The phylogenies of each NLP subfamilies followed closely the phylogenomic relationships of the genera in Dothideomycetes. Nevertheless, the imbalanced phylogeny of the NLP superfamily across the Dothideomycetes revealed that subfamilies underwent independent evolutionary paths, each of which showing different signatures of ancient gene duplications and biased successive gene losses that may be associated with changes in cytotoxic activity.

## Author contributions

All authors contributed to the study conception. TCSD performed data assembly and analyses. TCSD and LOO wrote the manuscript. HVSR commented and edited the manuscript. LOO and HVSR supervised the research. All authors read and approved the final manuscript.

## Additional files

**Additional file 1.** Detailed information on all fungal species used in this study.

**Additional file 2.** Detailed information on all NLPs in Fig. 2

